# The emergence of the rice blast fungus (*Pyricularia oryzae*) coincides with the evolutionary divergence of rice

**DOI:** 10.64898/2026.08.24.746645

**Authors:** Shunsuke Nozawa, Kazumichi Fujii

**Affiliations:** Fukushima Institute for Research, Education and Innovation, Namie, Fukushima, Japan; School of Agriculture, Tamagawa University, Machida, Tokyo, Japan

## Abstract

Rice blast, caused by *Pyricularia oryzae*, is one of the most destructive diseases threatening global rice production. Reconstructing the evolutionary history of *P. oryzae* in the context of the well-documented history of rice dispersal provides a powerful framework for predicting the coevolutionary dynamics of crops and their pathogens. However, the absence of fossil evidence has precluded robust estimation of the pathogen’s origin and evolutionary timescale. Here, we inferred genome-wide divergence times for *P. oryzae* using archaeological constraints derived from the emergence and spread of rice cultivation. Our analyses indicate that *P. oryzae* originated approximately 18,300 years ago, coincident with the divergence of *Oryza sativa* L. from its ancestral lineage rather than with rice domestication itself. Calibrating the evolutionary history of the pathogen against this host divergence further indicates that *P. oryzae* reached the Japanese archipelago 2,590 years ago based on nucleotide sequences and 3,030 years ago based on amino acid sequences, closely matching archaeological estimates for the arrival of rice agriculture in Japan. Contrary to the prevailing view that rice blast emerged with rice domestication approximately 10,000 years ago, our results place the origin of *P. oryzae* before domestication and link its subsequent spread to the expansion of cultivated rice. These findings redefine the evolutionary history of one of the world’s most important crop pathogens and establish a temporal framework for understanding crop–pathogen coevolution over millennial timescales.

## Introduction

How crop domestication and breeding shape the diversification and genetic homogenization of plant pathogens remains a central question in plant pathology^1, 2^. *Pyricularia oryzae* Cavara, the causal agent of rice blast disease, is posing a major threat to global food security, because rice is the staple food for more than half of the global population^3, 4^. The rice–*P. oryzae* pathosystem has long served as a model for understanding fungal pathogenicity and plant–pathogen interactions^5^.

Recent genome-scale studies have revealed that *P. oryzae* harbors its greatest genetic diversity in China and that extensive human-mediated gene flow from Chinese populations has played a central role in shaping population structure across East Asia, including Japan, Korea, and Taiwan^3^. Reconstructing the evolutionary history and dispersal of *P. oryzae* in the context of the well-documented history of rice dispersal provides a powerful framework for predicting the coevolutionary dynamics of crops and their pathogens.

The emergence of the rice-infecting lineage of *P. oryzae* has previously been dated using tip-dating based on isolate sampling dates. Based on these estimates, it has been proposed that the rice-infecting lineage emerged through a host shift from ancestral populations infecting *Setaria* species and that the increase in host population density associated with rice domestication approximately 10,000 years ago enabled the successful establishment and persistence of the pathogen in rice populations^6–11^. However, although tip dating is a powerful approach for estimating divergence times in rapidly evolving organisms such as viruses and bacteria, the relatively slow evolutionary rate of filamentous fungi often provides insufficient temporal signal for reliable molecular clock estimation^12^. Consistent with this limitation, our analysis showed that tip-dating does not adequately reflect the molecular clock in *P. oryzae* (Supplementary Fig. S1). Therefore, an independent temporal calibration framework is required to reassess the evolutionary history of the rice blast fungus. Recent genomic analyses have shown that *Oryza sativa*, diverged from its wild ancestor, *O. rufipogon*, approximately 18,300 years ago, and that some lineages lost the resistance gene against *P. oryzae* during this process^13^. Because the host shift to rice must have occurred after the emergence of *O. sativa*, this divergence provides a biologically meaningful upper temporal constraint on the origin of the rice-infecting lineage. Host evolution therefore provides an alternative calibration framework for estimating pathogen divergence times without relying on fossil evidence.

In parallel, archaeological evidence indicates that rice cultivation reached the Japanese archipelago approximately 2,700–3,000 years ago^14–18^. Because Japan has remained geographically isolated from continental East Asia, rice blast populations introduced during the spread of rice cultivation is assumed to have evolved independently. Consequently, the divergence of Japanese *P. oryzae* populations provides an independent benchmark for evaluating the plausibility of inferred evolutionary timescales.

Here, we reconstruct the evolutionary history of *P. oryzae* by calibrating divergence times using the split between *O. sativa* and *O. rufipogon*, and compared the resulting chronology with the archaeological records of rice dispersal into Japan.

## Results

### Host divergence provides a temporal calibration framework

Previous tip-dating analyses based on sampling years suggested that the rice blast fungus *P. oryzae* emerged approximately 10,000 years ago, coinciding with the domestication of rice. However, our analyses using the larger dataset detected little temporal signal in sampling dates, indicating that they are unsuitable for reliable molecular-clock calibration and calling previous timescale estimates into question (Supplementary Fig. 1). This discrepancy between tip-dating estimates and the observed evolutionary signal highlights the need for an independent calibration framework to infer a more realistic evolutionary timescale.

To address this issue, we reconstructed genome-scale phylogenies using 91 genomes of *P. oryzae* isolates infecting *O. sativa*, together with isolates from the *Setaria* lineage, which has been proposed as the ancestral source of the host shift to rice. We reasoned that major divergence events in the pathogen were conservatively constrained by the evolutionary history of its host and therefore calibrated the pathogen phylogeny using the divergence between *O. sativa* and *O. rufipogon*, estimated at approximately 18,300 years before present^13^.

Phylogenetic analyses based on 2,766 single-copy orthologous genes (>5.5 Mb nucleotide and >1.8 Mb amino acid) produced congruent phylogenetic trees based on nucleotide and amino acid data on earliest Japanese lineage in the phylogeny (Supplementary figure 2 and 3). In both datasets, the rice-infecting lineage formed a sister clade to isolates infecting green foxtail (*Setaria viridis*), supporting a host shift from a *Setaria*-associated ancestor. Within the rice-infecting lineage, an isolate from Hubei, China is located the earliest-diverging position, whereas the remaining rice isolates formed a sister clade, reinforcing previous evidence for a Chinese origin of the rice blast pathogen^3^.

We further identified the earliest lineage associated with the introduction of rice blast into the Japanese archipelago (Figs. 2 and 3). Divergence dating based on nucleotide sequences place the event at approximately 2,590 years before present, and estimates based on amino acid sequences place it at approximately 3,030 years before present. Despite these differences, both analyses consistently indicate that the introduction of rice blast into Japan occurred during the period when cultivated rice first spread into the archipelago.

**Fig. 1.**
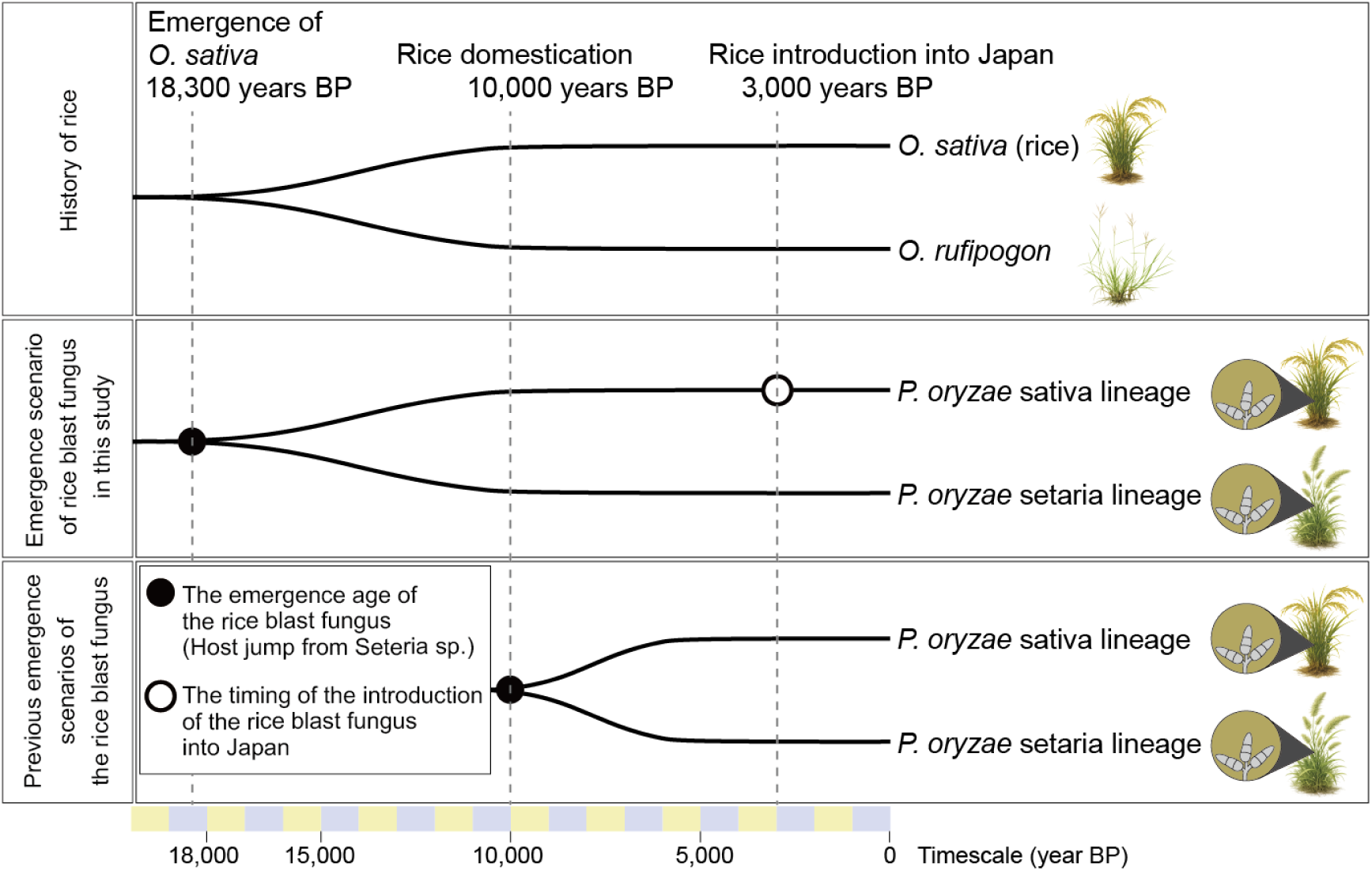
Evolutionary scenario of the rice blast fungus *Pyricularia oryzae*. This study tests the hypothesis that the divergence between *Oryza rufipogon* and *O. sativa* (18,300 years ago) coincides with the inferred host jump of *Pyricularia oryzae* from *Setaria* species. Using the introduction of rice agriculture into the Japanese archipelago (approximately 3,000 years BP) as a chronological anchor, we apply cross-dating approaches to evaluate the evolutionary timescale of the rice blast fungus and compare it with the prevailing view that the pathogen emerged around 10,000 years ago during rice domestication.

**Fig. 2.**
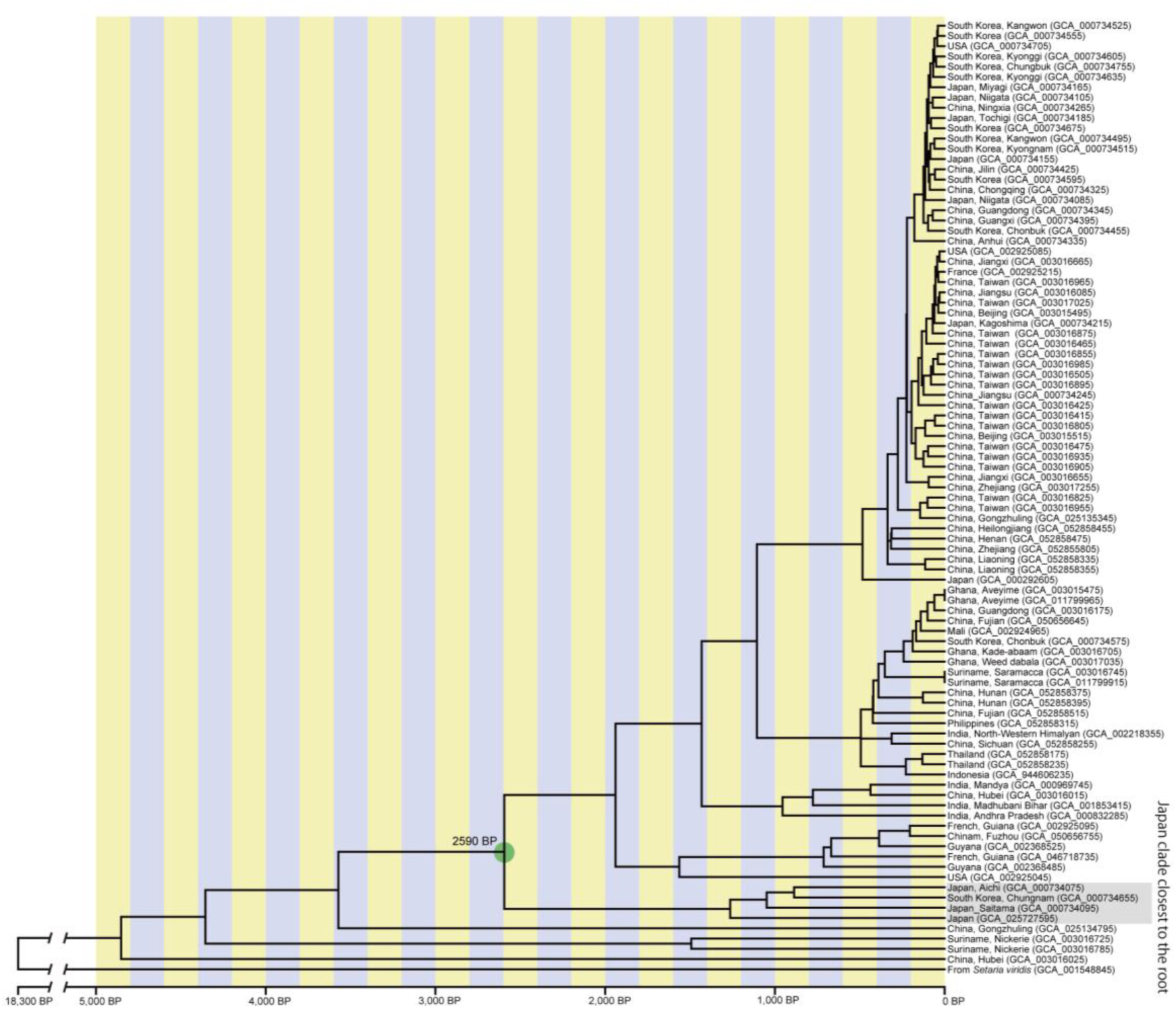
Host-calibrated, time-scaled phylogeny of *Pyricularia oryzae* based on nucleotide sequence data showing consistency with the archaeological timing of rice dispersal into the Japanese archipelago. A strict molecular-clock phylogeny calibrated using the host divergence between *Oryza sativa* and *O. rufipogon* (18,300 BP) places the divergence from the oldest Japanese lineage at 2,590 BP, consistent with the archaeological timing of rice dispersal into the Japanese archipelago.

**Fig. 3.**
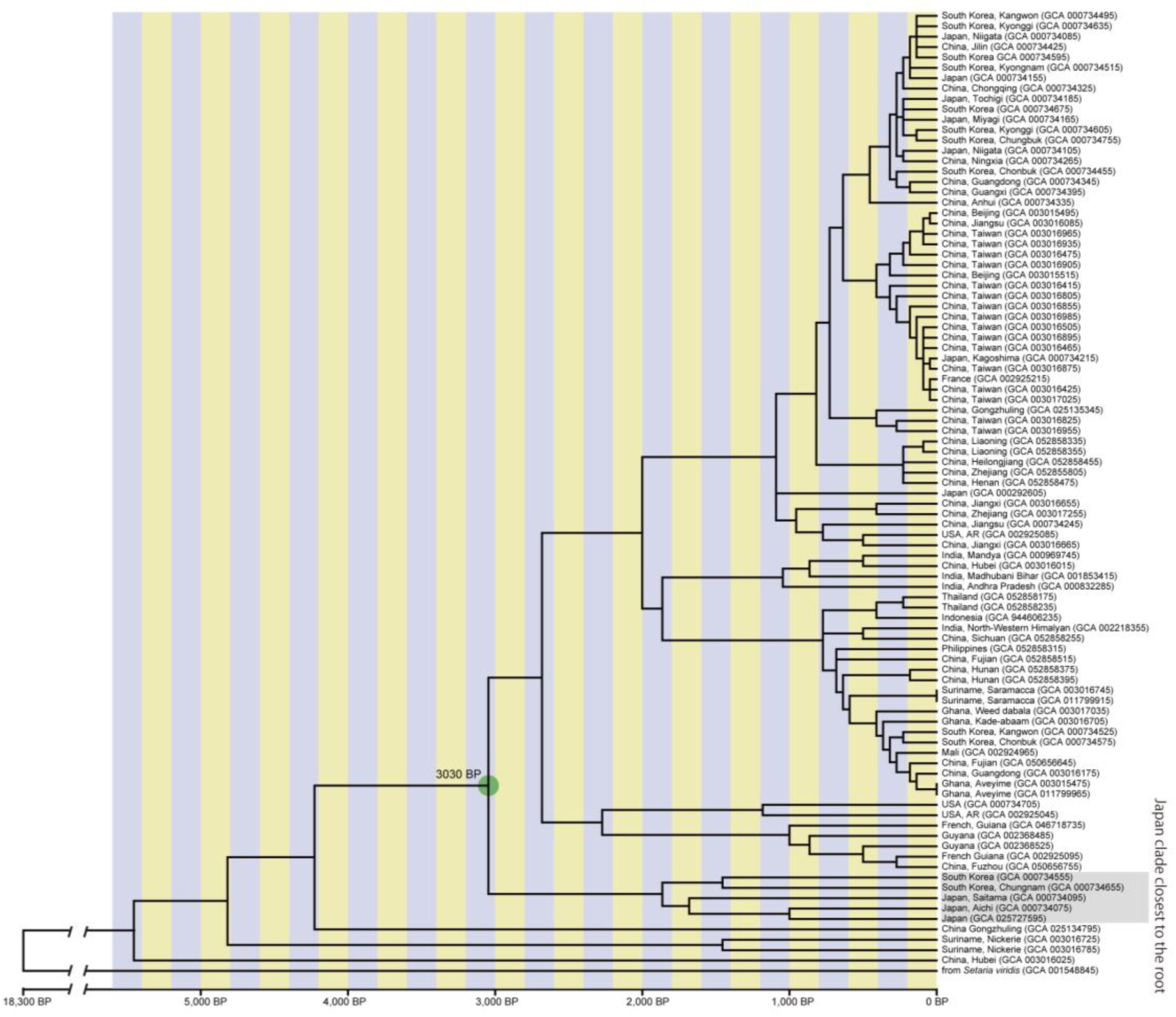
Host-calibrated, time-scaled phylogeny of *Pyricularia oryzae* based on amino acid sequence data showing consistency with the archaeological timing of rice dispersal into the Japanese archipelago. A strict molecular-clock phylogeny calibrated using the host divergence between *Oryza sativa* and *O. rufipogon* (18,300 BP) places the divergence from the oldest Japanese lineage at 3,030 BP, consistent with the archaeological timing of rice dispersal into the Japanese archipelago.

### Molecular dating links the introduction of rice blast to the spread of rice cultivation

Archaeological evidence indicates that rice agriculture spread from continental East Asia into northern Kyushu approximately 3,000–2,700 years ago and subsequently expanded throughout western Japan. The close correspondence between these archaeological estimates and our host-calibrated molecular dating supports the hypothesis that the rice-infecting lineage of *P. oryzae* emerged shortly after the divergence of *O. sativa* and *O. rufipogon* (approximately 18,300 years before present)^12^ (Choi et al., 2017) and was subsequently introduced into the Japanese archipelago together with cultivated rice, where it became established as rice agriculture spread (Fig. 4). Thus, pathogen phylogenies may provide an independent temporal framework that complements archaeological evidence for prehistoric crop dispersal.

Not all pathogen lineages, however, retained evolutionary signatures consistent with the historical dispersal of rice. Present-day populations have likely been shaped by lineage extinction, lineage replacement, incomplete sampling of ancestral diversity, and extensive human-mediated dispersal over the past several millennia. For example, rice cultivation in Suriname originated only in the late seventeenth century, followed by repeated introductions of rice varieties from India, China, and Java during the nineteenth and twentieth centuries. Consequently, Surinamese isolates that diverged earlier than the oldest Japanese lineage identified here are unlikely to represent relics of ancient dispersal events. Instead, they most likely reflect more recent introductions from unsampled Asian populations, particularly those from China. These findings highlight the limitations of reconstructing ancient migration histories solely from extant pathogen populations.

Our estimated timescale differs substantially from the prevailing hypothesis that the rice-infecting lineage emerged during rice domestication approximately 10,000 years ago^11^. Instead, our results suggest that the rice-infecting lineage originated around the time of divergence between *O. sativa* and *O. rufipogon*. Previous studies have proposed that the rice blast pathogen arose through a host jump from a *Setaria*-infecting ancestor. Because green foxtail (*S. viridis*) diverged earlier than the genus *Oryza* and likely co-occurred with ancestral or early domesticated rice populations, ecological overlap between these hosts may have facilitated pathogen exchange and host shifts. Our estimated divergence times are therefore consistent with a scenario in which the emergence of the rice-infecting lineage accompanied the early evolutionary history of rice.

Taken together, a host-calibrated phylogenomic framework estimated the introduction of rice blast into Japan at 2,590 or 3,030 years before present, in close agreement with archaeological evidence for expansion of rice cultivation. These findings suggest that the evolutionary history of the rice blast fungus is more closely linked to host divergence and subsequent crop dispersal than previously recognized, providing a revised scenario for understanding the origin and spread of one of the world’s most important plant pathogens.

## Conclusion

Here, we showed that (1) the rice blast fungus population lacks the temporal signal required for reliable tip-dating analyses, (2) a host-calibrated phylogeny based on the divergence between *O. sativa* and *O. rufipogon* dated the earliest introduction of the rice blast fungus into Japan to approximately 2,590 or 3,030 years before present, and (3) this estimate is consistent with archaeological evidence for the spread of rice cultivation into Japan, as the earliest archaeologically documented introduction of rice agriculture to the Japanese archipelago is generally dated to approximately 2,700–3,000 years before present. Together, these results indicate that the emergence of the rice-infecting lineage was more closely associated with host evolution than with the domestication of rice itself.

The evolutionary timescale presented here is based on the calibration hypothesis that the divergence between the rice-infecting lineage and its closest *Setaria*-infecting lineage corresponds to the divergence between *O. sativa* and *O. rufipogon*. Furthermore, given the likely ecological overlap between broadly distributed *Setaria* species and ancestral rice populations, our estimated divergence time is consistent with an evolutionary scenario in which the rice blast fungus arose through a host shift.

Our analyses are limited by incomplete geographical sampling, which remains insufficient to fully reconstruct the dispersal routes and population expansion of the rice blast fungus into the Japanese archipelago. Future studies incorporating additional genome sequences from inland China, the Korean Peninsula, Southeast Asia, and other underrepresented regions, together with the host-calibrated framework presented here, will enable a more comprehensive reconstruction of the evolutionary history and regional diversification of this pathogen.

Taken together, our study revises the evolutionary history of the rice blast fungus by proposing an alternative evolutionary scenario to the prevailing hypothesis that the rice-infecting lineage emerged only after rice domestication. More broadly, our study demonstrates the utility of host evolutionary history as an independent temporal framework for dating pathogen evolution when conventional molecular-clock approaches are limited. Integrating comparative genomics, host evolution, and archaeological evidence provides a general framework for reconstructing the origins, dispersal and diversification of plant pathogens and their co-evolution with agriculture.

## Materials and methods

### Taxon Sampling

Genome data were obtained from the NCBI database, including the fungal isolates used in Duan et al.^3^ and Zhong et al.^11^. To determine the regional origins of Japanese fungal isolates, we prioritized genomes from samples that provided detailed locality information, particularly those specifying Chinese provinces and Japanese prefectures (Supplementary Table 1). The ingroup taxa comprised the 94 genomic data of *Pyricularia oryzae* isolated from *Oryza sativa* originated from China, Cote d’Ivoire, France, French Guiana, Ghana, Guyana, India, Indonesia, Japan, Mali, Philippines, South Korea, Suriname, Thailand, and USA. As outgroup taxa, one strain of *P. oryzae* isolated from *Setaria viridis* and one strain of *P. grisea*, and *P. penniseti* were used. In particular, *P. oryzae* isolates obtained from *S. viridis* were used to define the divergence point from rice-infecting lineages, and this divergence was incorporated into the temporal calibration of the phylogeny.

### Orthology Inference of Genes and Selection

Genes were predicted from genome data using Augustus v.3.3.3^19^ with the parameter: “--genemodel = complete --species = Magnaporthe grisea <GENOMIC data>”. OrthoFinder v2.5.4^20^ was used to identify single-copy orthologs shared among taxa using default parameters (inflation parameter 1.5). Obtained single-copy orthologs were aligned with MAFFT v7.505^21^ with default setting. Gaps including sites in alignments were removed using trimAl v1.415^22^ with the command option “-g 10”. Sequence statistics for the concatenated datasets were calculated using the Sequence Data Explorer module of MEGA10^23^.

### Phylogenetic Analyses

Phylogenetic trees were reconstructed using the concatenation approach. Concatenated alignments of nucleotide and amino acid sequences from all 2,766 single-copy orthologous genes (Supplementary Table 2) were analyzed separately using IQ-TREE v.7.0.4^24^. The nucleotide alignment was analyzed under the GTR+G model, whereas the amino acid alignment was analyzed under the LG+G model, with 1,000 bootstrap replicates for each analysis.

### Evaluation of temporal signal for tip-dating

To assess whether tip-dating was appropriate for estimating divergence times in *P. oryzae*, the temporal signal contained within the genomic dataset was evaluated. A maximum-likelihood phylogeny inferred from whole-genome SNP data was rooted using *P. grisea* and *P. penniseti* as outgroups. Root-to-tip distances were calculated for all isolates using the R package ape, and their relationship with sampling year was examined by linear regression.

To determine whether the observed association exceeded that expected by chance, a date randomization test (DRT) was performed. Sampling years were randomly reassigned among isolates 1,000 times while maintaining the original phylogeny. For each randomized dataset, root-to-tip regression was repeated and the coefficient of determination (*R*²) was recorded. The observed *R*² value was compared with the distribution obtained from randomized datasets. Because tip-dating assumes a positive relationship between sampling time and genetic divergence, both the strength and direction of the regression slope were evaluated.

Because tip-dating assumes that genetic divergence accumulates through time, the presence of temporal signal was evaluated prior to any interpretation of divergence-time estimates.

### Divergence time estimation

Divergence times were estimated using the RelTime method implemented in MEGA-CC (Molecular Evolutionary Genetics Analysis Command Line). The maximum-likelihood phylogeny inferred from the concatenated alignment of 2,766 single-copy orthologous genes was used as the input tree. The tree was rooted using *P. grisea* and *P. penniseti* as outgroups.

Because no significant temporal signal was detected among the sampling dates (Supplementary figure 1), tip-dating was not applied. Instead, divergence time estimation was calibrated using the divergence between the *Setaria viridis*-infecting lineage and the rice-infecting lineage of *P. oryzae*. This node was fixed at 18,300 years before present, corresponding to the estimated divergence time between domesticated rice (*O. sativa*) and its wild progenitor (*O. rufipogon*) reported by Choi et al.^13^. Divergence times for all remaining nodes were subsequently estimated using the RelTime algorithm implemented in MEGA-CC under the default settings. The same procedure was independently applied to the amino acid alignment to estimate divergence times based on the amino acid phylogeny.

## Acknowledgments

We thank Professor Kyoko Watanabe (Tamagawa University) for her helpful advice and insightful comments on this work.

## Data Availability

The datasets in this study are available in Supplementary data.

## Author Contribution

S.N. conceived the study, designed the methodology, conducted all analyses, visualized the data, and led manuscript preparation. K. F. assisted in designing the methodology and manuscript editing. All authors have reviewed and approved the final manuscript.

## Competing Interests Statement

The authors declare no competing interests.

---

**Supplementary Fig. 1.**
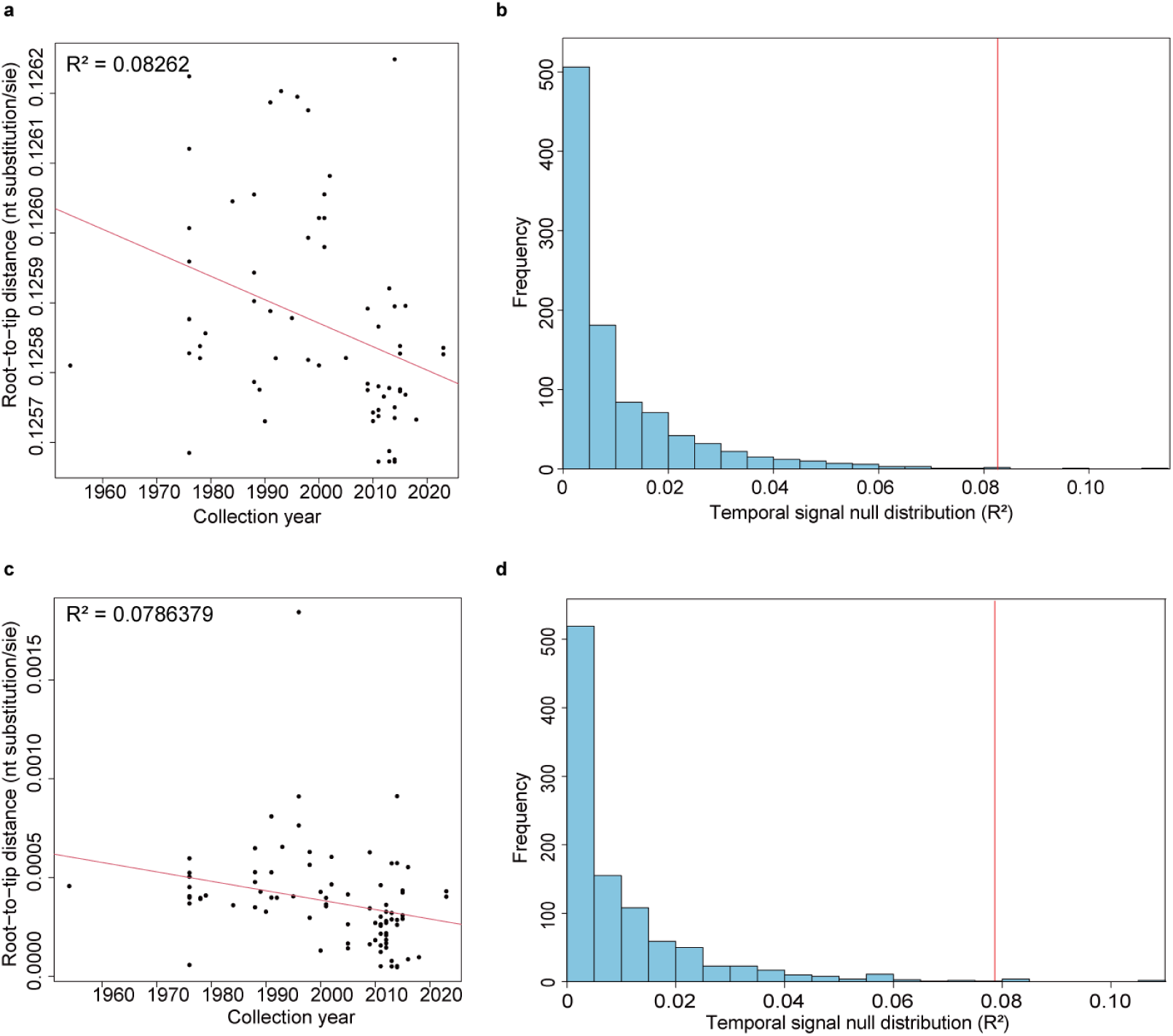
Assessment of temporal signal in the dataset. a, Root-to-tip regression based on nucleotide data showing a negative relationship between genetic divergence and collection year with low a R² value (0.0826), indicating that more recent isolates tend to fall closer to the root. b, Date-randomization test (1,000 permutations) demonstrating that randomized R² values cluster near zero, with a 95% upper bound of 0.048. The observed R² (0.0826) falls well within the null distribution (95% upper bound = 0.121), indicating that the dataset lacks meaningful temporal signal and does not meet the assumptions required for reliable tip-dating. c, Consistent with the result of nucleotide data, root-to-tip regression based on amino-acid sequence, showing a negative relationship between genetic divergence and collection year with low a R² value (0.0786). d, Date-randomization test (1,000 permutations) demonstrating that the observed R² (0.0786) falls well within the null distribution (95% upper bound = 0.109), indicating that the dataset lacks meaningful temporal signal and does not meet the assumptions required for reliable tip-dating. Together, these results demonstrate that neither nucleotide nor amino-acid datasets provide insufficient clock-like structure for robust tip-dating analyses.

**Supplementary Fig. 2.**
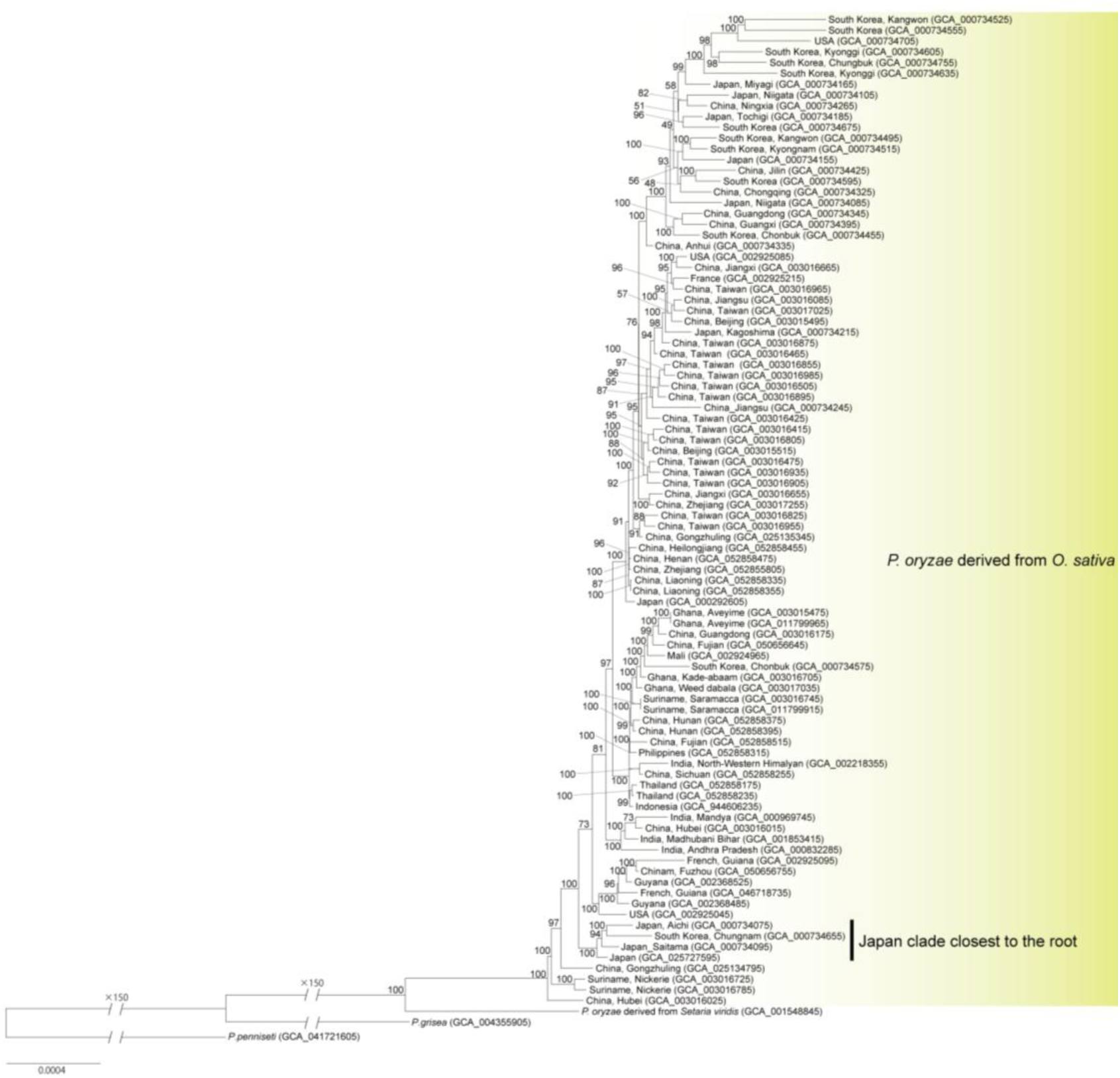
Global phylogeny of *Pyricularia oryzae* based on nucleotide sequence data. Maximum-likelihood phylogeny constructed from a combined dataset of 2,766 single copy genes shows the Chinese isolates occupied the deepest position within the *O. sativa*–derived lineage, and one clade consists of Japanese isolates including one Korean isolate is distinct from other isolates. Outgroups (*P. grisea*, *P. pennisetii* and *P. oryzae* from *Setaria viridis*) confirm the basal placement of all the *O. sativa*–derived *P. oryzae* lineage. Scale bar represents 0.0004 substitutions per site. Bootstrap values are shown at each node

**Supplementary Fig. 3.**
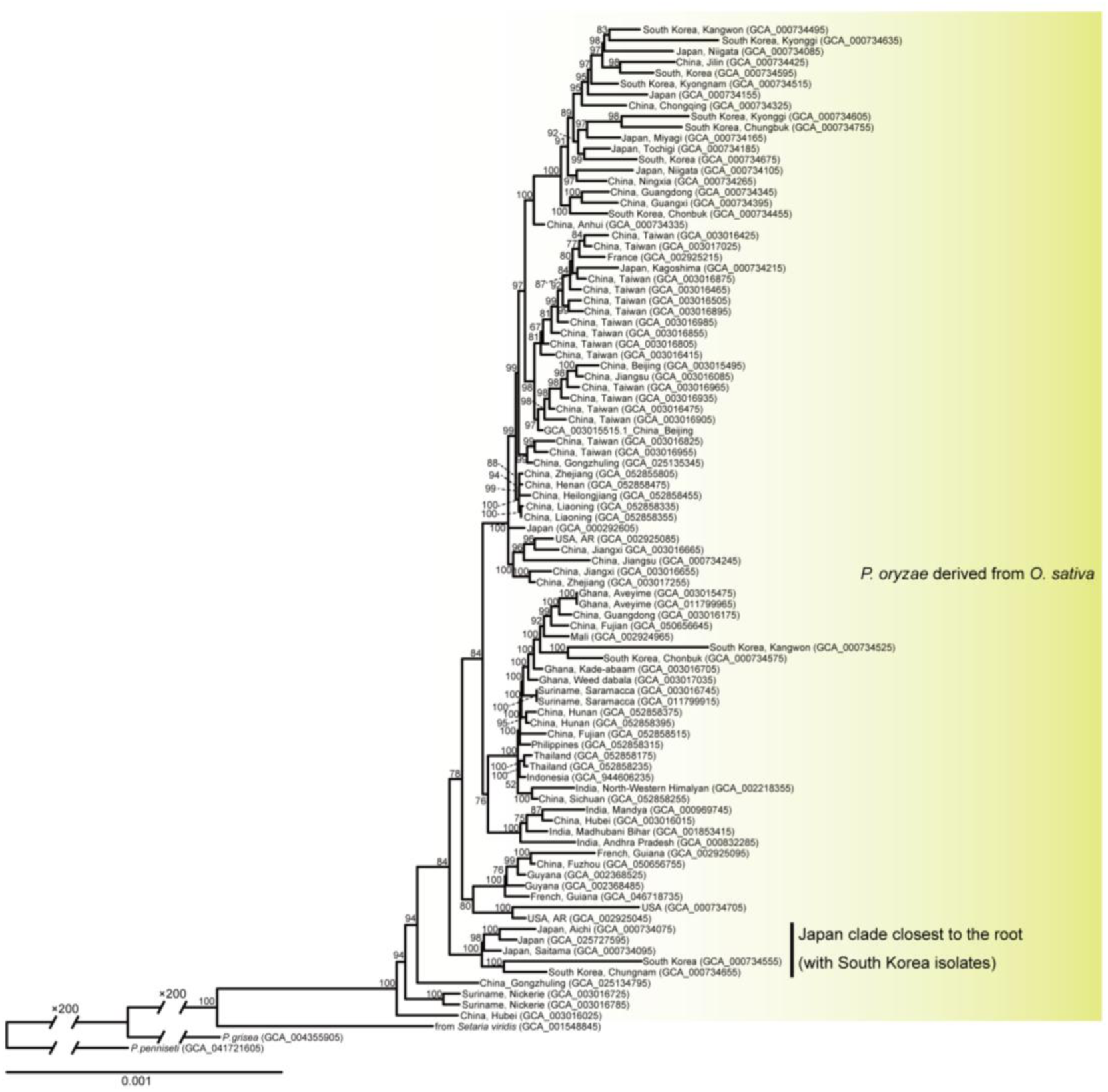
Global phylogeny of *Pyricularia oryzae* based on amino acid sequence data. Maximum-likelihood phylogeny constructed from a combined dataset of 2,766 single copy genes shows the Chinese isolates occupied the deepest position within the *O. sativa*–derived lineage, and one clade consists of Japanese isolates including one Korean isolate is distinct from other isolates. Outgroups (*P. grisea*, *P. pennisetii* and *P. oryzae* from *Setaria viridis*) confirm the basal placement of all the *O. sativa*–derived *P. oryzae* lineage. Scale bar represents 0.001 substitutions per site. Bootstrap values are shown at each node

**Supplementary Table 1.**
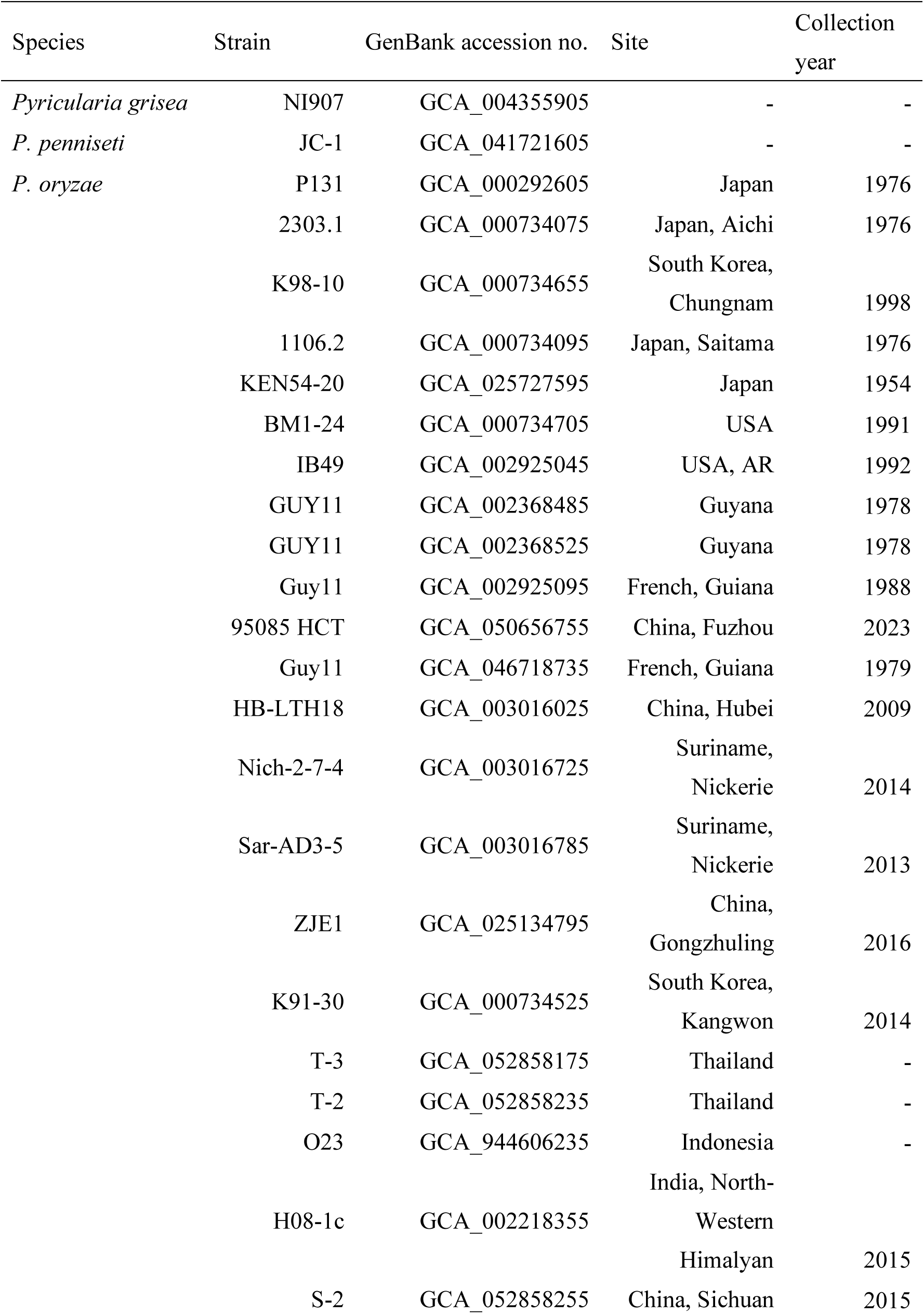
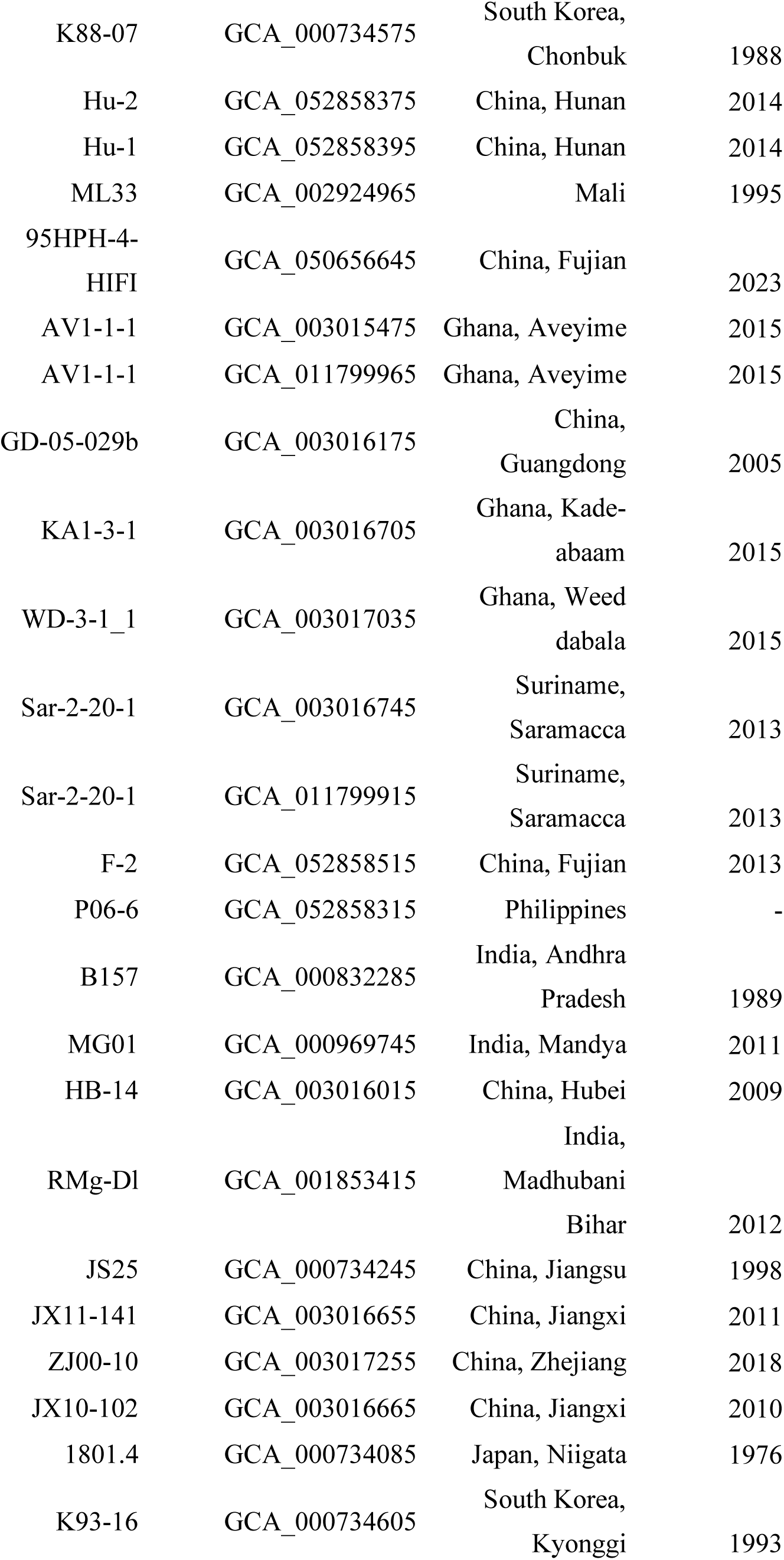
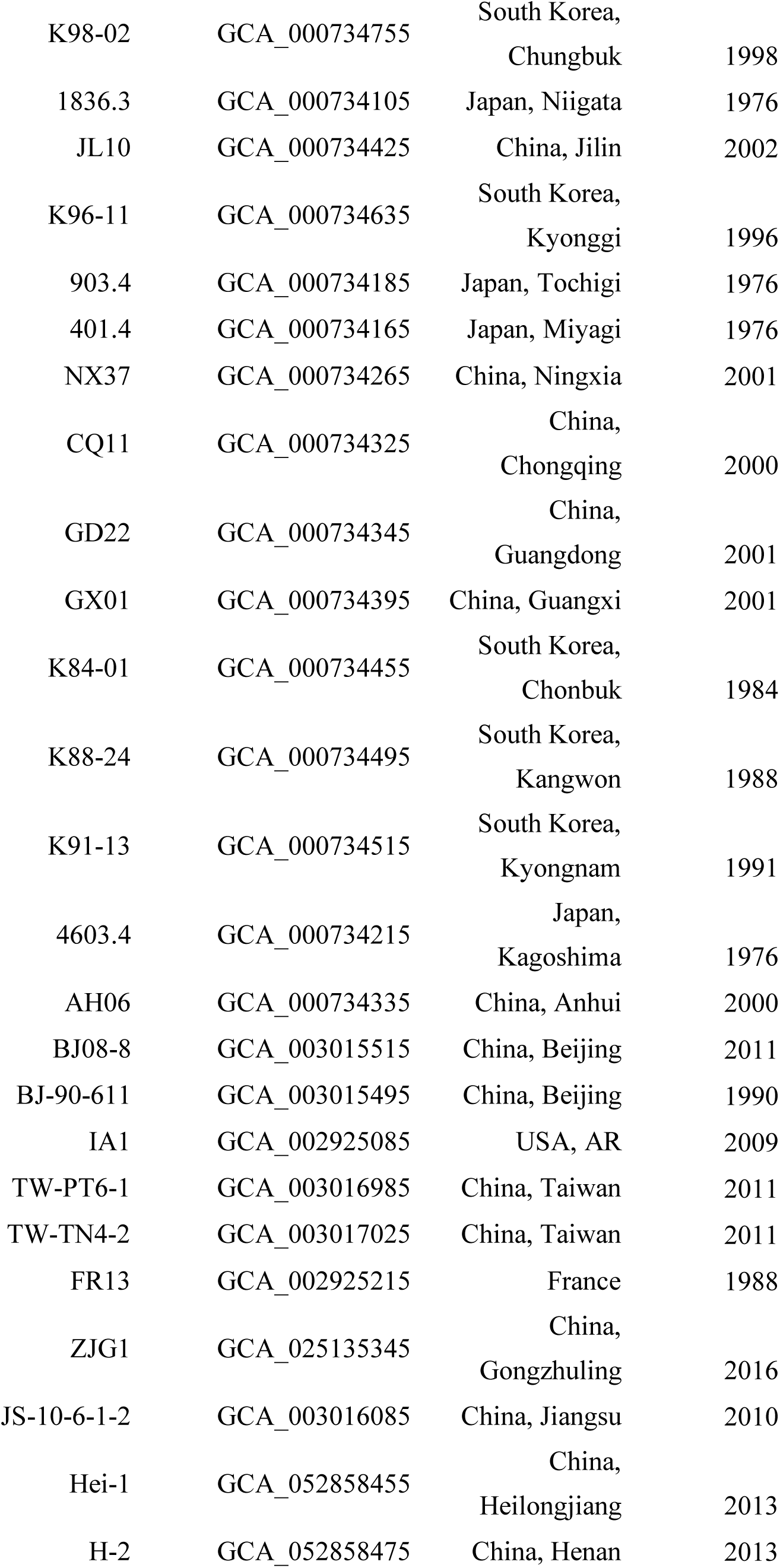
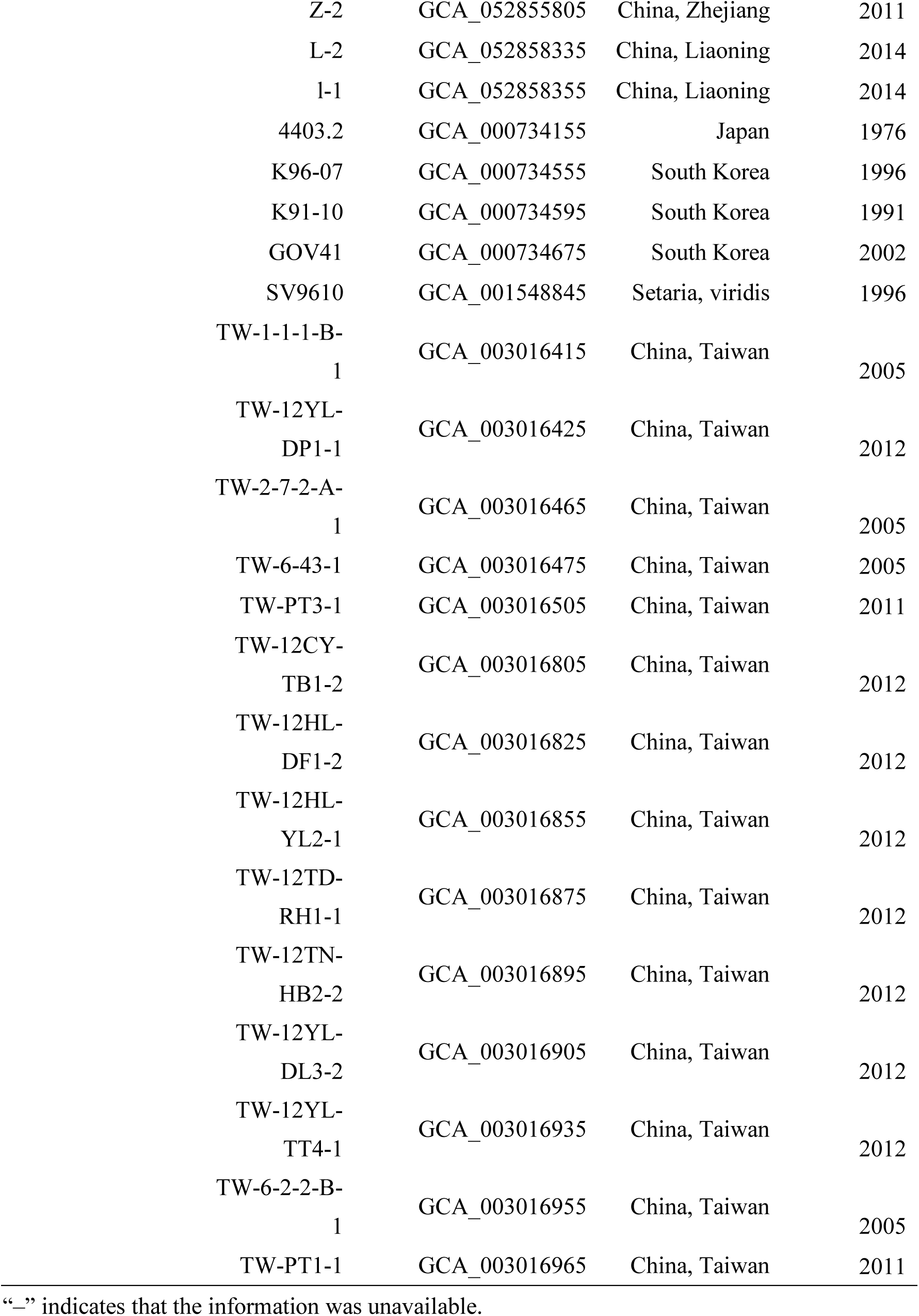
Information on isolates used in this study.

| Species | Strain | GenBank accession no. | Site | Collection year |
| --- | --- | --- | --- | --- |
| *Pyricularia grisea* | NI907 | GCA_004355905 | - | - |
| *P. penniseti* | JC-1 | GCA_041721605 | - | - |
| *P. oryzae* | P131 | GCA_000292605 | Japan | 1976 |
|  | 2303.1 | GCA_000734075 | Japan, Aichi | 1976 |
|  | K98-10 | GCA_000734655 | South Korea, Chungnam | 1998 |
|  | 1106.2 | GCA_000734095 | Japan, Saitama | 1976 |
|  | KEN54-20 | GCA_025727595 | Japan | 1954 |
|  | BM1-24 | GCA_000734705 | USA | 1991 |
|  | IB49 | GCA_002925045 | USA, AR | 1992 |
|  | GUY11 | GCA_002368485 | Guyana | 1978 |
|  | GUY11 | GCA_002368525 | Guyana | 1978 |
|  | Guy11 | GCA_002925095 | French, Guiana | 1988 |
|  | 95085 HCT | GCA_050656755 | China, Fuzhou | 2023 |
|  | Guy11 | GCA_046718735 | French, Guiana | 1979 |
|  | HB-LTH18 | GCA_003016025 | China, Hubei | 2009 |
|  | Nich-2-7-4 | GCA_003016725 | Suriname, Nickerie | 2014 |
|  | Sar-AD3-5 | GCA_003016785 | Suriname, Nickerie | 2013 |
|  | ZJE1 | GCA_025134795 | China, Gongzhuling | 2016 |
|  | K91-30 | GCA_000734525 | South Korea, Kangwon | 2014 |
|  | T-3 | GCA_052858175 | Thailand | - |
|  | T-2 | GCA_052858235 | Thailand | - |
|  | O23 | GCA_944606235 | Indonesia | - |
|  | H08-1c | GCA_002218355 | India, North-Western Himalyan | 2015 |
|  | S-2 | GCA_052858255 | China, Sichuan | 2015 |

|  |  |  |  |
| --- | --- | --- | --- |
| K88-07 | GCA_000734575 | South Korea,<br>Chonbuk | 1988 |
| Hu-2 | GCA_052858375 | China, Hunan | 2014 |
| Hu-1 | GCA_052858395 | China, Hunan | 2014 |
| ML33 | GCA_002924965 | Mali | 1995 |
| 95HPH-4-HIFI | GCA_050656645 | China, Fujian | 2023 |
| AV1-1-1 | GCA_003015475 | Ghana, Aveyime | 2015 |
| AV1-1-1 | GCA_011799965 | Ghana, Aveyime | 2015 |
| GD-05-029b | GCA_003016175 | China,<br>Guangdong | 2005 |
| KA1-3-1 | GCA_003016705 | Ghana, Kade-<br>abaam | 2015 |
| WD-3-1_1 | GCA_003017035 | Ghana, Weed<br>dabala | 2015 |
| Sar-2-20-1 | GCA_003016745 | Suriname,<br>Saramacca | 2013 |
| Sar-2-20-1 | GCA_011799915 | Suriname,<br>Saramacca | 2013 |
| F-2 | GCA_052858515 | China, Fujian | 2013 |
| P06-6 | GCA_052858315 | Philippines | - |
| B157 | GCA_000832285 | India, Andhra<br>Pradesh | 1989 |
| MG01 | GCA_000969745 | India, Mandya | 2011 |
| HB-14 | GCA_003016015 | China, Hubei | 2009 |
| RMg-DI | GCA_001853415 | India,<br>Madhubani<br>Bihar | 2012 |
| JS25 | GCA_000734245 | China, Jiangsu | 1998 |
| JX11-141 | GCA_003016655 | China, Jiangxi | 2011 |
| ZJ00-10 | GCA_003017255 | China, Zhejiang | 2018 |
| JX10-102 | GCA_003016665 | China, Jiangxi | 2010 |
| 1801.4 | GCA_000734085 | Japan, Niigata | 1976 |
| K93-16 | GCA_000734605 | South Korea,<br>Kyonggi | 1993 |
| K98-02 | GCA_000734755 | South Korea,<br>Chungbuk | 1998 |
| 1836.3 | GCA_000734105 | Japan, Niigata | 1976 |
| JL10 | GCA_000734425 | China, Jilin | 2002 |
| K96-11 | GCA_000734635 | South Korea,<br>Kyonggi | 1996 |
| 903.4 | GCA_000734185 | Japan, Tochigi | 1976 |
| 401.4 | GCA_000734165 | Japan, Miyagi | 1976 |
| NX37 | GCA_000734265 | China, Ningxia | 2001 |
| CQ11 | GCA_000734325 | China,<br>Chongqing | 2000 |
| GD22 | GCA_000734345 | China,<br>Guangdong | 2001 |
| GX01 | GCA_000734395 | China, Guangxi | 2001 |
| K84-01 | GCA_000734455 | South Korea,<br>Chonbuk | 1984 |
| K88-24 | GCA_000734495 | South Korea,<br>Kangwon | 1988 |
| K91-13 | GCA_000734515 | South Korea,<br>Kyongnam | 1991 |
| 4603.4 | GCA_000734215 | Japan,<br>Kagoshima | 1976 |
| AH06 | GCA_000734335 | China, Anhui | 2000 |
| BJ08-8 | GCA_003015515 | China, Beijing | 2011 |
| BJ-90-611 | GCA_003015495 | China, Beijing | 1990 |
| IA1 | GCA_002925085 | USA, AR | 2009 |
| TW-PT6-1 | GCA_003016985 | China, Taiwan | 2011 |
| TW-TN4-2 | GCA_003017025 | China, Taiwan | 2011 |
| FR13 | GCA_002925215 | France | 1988 |
| ZJG1 | GCA_025135345 | China,<br>Gongzhuling | 2016 |
| JS-10-6-1-2 | GCA_003016085 | China, Jiangsu | 2010 |
| Hei-1 | GCA_052858455 | China,<br>Heilongjiang | 2013 |
| H-2 | GCA_052858475 | China, Henan | 2013 |
| Z-2 | GCA_052855805 | China, Zhejiang | 2011 |
| L-2 | GCA_052858335 | China, Liaoning | 2014 |
| l-1 | GCA_052858355 | China, Liaoning | 2014 |
| 4403.2 | GCA_000734155 | Japan | 1976 |
| K96-07 | GCA_000734555 | South Korea | 1996 |
| K91-10 | GCA_000734595 | South Korea | 1991 |
| GOV41 | GCA_000734675 | South Korea | 2002 |
| SV9610 | GCA_001548845 | *Setaria, viridis* | 1996 |
| TW-1-1-1-B-1 | GCA_003016415 | China, Taiwan | 2005 |
| TW-12YL-DP1-1 | GCA_003016425 | China, Taiwan | 2012 |
| TW-2-7-2-A-1 | GCA_003016465 | China, Taiwan | 2005 |
| TW-6-43-1 | GCA_003016475 | China, Taiwan | 2005 |
| TW-PT3-1 | GCA_003016505 | China, Taiwan | 2011 |
| TW-12CY-TB1-2 | GCA_003016805 | China, Taiwan | 2012 |
| TW-12HL-DF1-2 | GCA_003016825 | China, Taiwan | 2012 |
| TW-12HL-YL2-1 | GCA_003016855 | China, Taiwan | 2012 |
| TW-12TD-RH1-1 | GCA_003016875 | China, Taiwan | 2012 |
| TW-12TN-HB2-2 | GCA_003016895 | China, Taiwan | 2012 |
| TW-12YL-DL3-2 | GCA_003016905 | China, Taiwan | 2012 |
| TW-12YL-TT4-1 | GCA_003016935 | China, Taiwan | 2012 |
| TW-6-2-2-B-1 | GCA_003016955 | China, Taiwan | 2005 |
| TW-PT1-1 | GCA_003016965 | China, Taiwan | 2011 |

“—” indicates that the information was unavailable.

**Supplementary Table 2.**
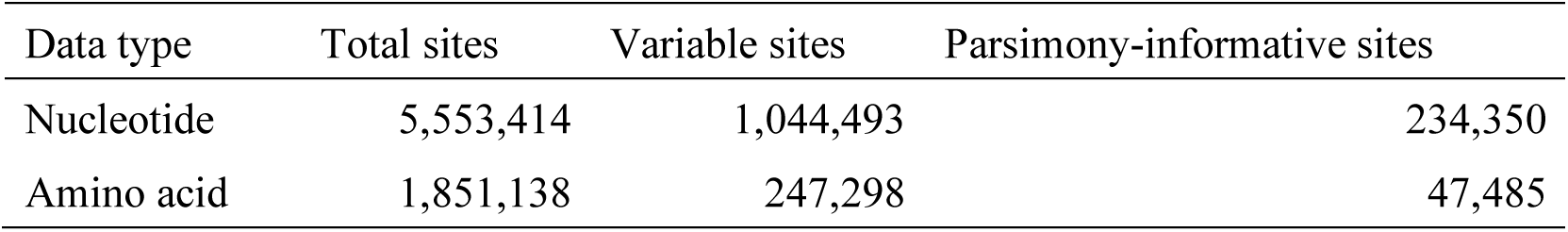
Summary statistics of concatenated superalignment datasets from 2,766 single-copy orthologous.

## Notes

### Competing Interest Statement

The authors have declared no competing interest.

## Reference

1. Madhushan, A. et al. From natural hosts to agricultural threats: The evolutionary journey of phytopathogenic fungi. Journal of Fungi, 11, 25 (2025).

2. Feurtey, A. et al. A thousand-genome panel retraces the global spread and adaptation of a major fungal crop pathogen. Nature communications 14, 1059 (2023).

3. Duan, G. et al. Chinese populations of Magnaporthe oryzae serving as a source of human-mediated gene flow to Asian countries: A population genomic analysis. Journal of Fungi, 10, 739 (2024).

4. Bandumula, N. Rice production in Asia: Key to global food security. Proc. Natl Acad. Sci. India Sect. B Biol. Sci. 88, 1323–1328 (2018).

5. Dean, R. et al. The Top 10 fungal pathogens in molecular plant pathology. Mol. Plant Pathol 13, 414–430 (2012).

6. Gladieux, P. et al. Gene flow between divergent cereal- and grass-specific lineages of the rice blast fungus Magnaporthe oryzae. mBio 9, e01219–18 (2018).

7. Crawford, G. W., Chen, X. & Wang, J. Houli culture rice from the Yuezhuang site, Jinan. Dongfang Kaogu 3, 247–251 (2006).

8. Fuller, D. Q. et al. The domestication process and domestication rate in rice: spikelet bases from the Lower Yangtze. Science 323, 1607–1610 (2009).

9. Zhao, Z. New archaeobotanic data for the study of the origins of agriculture in China. Curr. Anthropol. 52, S295–S306 (2011).

10. Couch, B. C. et al. Origins of host-specific populations of the blast pathogen Magnaporthe oryzae in crop domestication with subsequent expansion of pandemic clones on rice and weeds of rice. Genetics 170, 613–630 (2005).

11. Zhong, Z. et al. Population genomic analysis of the rice blast fungus reveals specific events associated with expansion of three main clades. ISME J. 12, 1867–1878 (2018).

12. Edwards, H. M. & Rhodes, J. Accounting for the biological complexity of pathogenic fungi in phylogenetic dating. J. Fungi 7, 661 (2021).

13. Choi, J. Y. et al. The rice paradox: multiple origins but single domestication in Asian rice. Mol. Biol. Evol. 34, 969–979 (2017).

14. Leipe, C. et al. The spread of rice to Japan: insights from Bayesian analysis of direct radiocarbon dates and population dynamics in East Asia. Quat. Sci. Rev. 244, 106507 (2020).

15. Miyamoto, K. The spread of rice agriculture during the Yayoi period: From the Shandong Peninsula to Japanese Archipelago via Korean Peninsula. Jpn. J. Archaeol. 6, 109–124 (2019).

16. Toyama, S. The beginning of rice cultivation and changes in land conditions as identified by plant opal analysis. Quat. Res. 33, 317–329 (1994).

17. Craig, O. E., et al. Lipid residue analysis reveals divergent culinary practices in Japan and Korea at the dawn of intensive agriculture. Proc. Natl Acad. Sci. USA 122, e2504414122 (2025).

18. Crema, E. R., Stevens, C. J. & Shoda, S. Bayesian analyses of direct radiocarbon dates reveal geographic variations in the rate of rice farming dispersal in prehistoric Japan. Sci. Adv. 8, eadc9171 (2022).

19. Stanke, M. & Morgenstern, B. AUGUSTUS: a web server for gene prediction in eukaryotes that allows user-defined constraints. Nucleic Acids Res. 33, W465–W467 (2005).

20. Emms, D. M. & Kelly, S. OrthoFinder: phylogenetic orthology inference for comparative genomics. Genome Biol. 20, 238 (2019).

21. Katoh, K. & Standley, D. M. MAFFT multiple sequence alignment software version 7: improvements in performance and usability. Mol. Biol. Evol. 30, 772–780 (2013).

22. Capella-Gutiérrez, S., Silla-Martínez, J. M. & Gabaldón, T. trimAl: a tool for automated alignment trimming in large-scale phylogenetic analyses. Bioinformatics 25, 1972–1973 (2009).

23. Kumar, S. et al. MEGA X: Molecular evolutionary genetics analysis across computing platforms. Mol. Biol. Evol. 35, 1547–1549 (2018).

24. Stamatakis, A. RAxML version 8: a tool for phylogenetic analysis and post-analysis of large phylogenies. Bioinformatics 30, 1312–1313 (2014).

